# Neurons define non-myelinated axon segments by the regulation of galectin-4-containing axon membrane domains

**DOI:** 10.1101/115758

**Authors:** Natalia Díez-Revuelta, Alonso M. Higuero, Silvia Velasco, María Peñas-de-la-Iglesia, Hans-Joachim Gabius, José Abad-Rodríguez

**Affiliations:** Membrane Biology and Axonal Repair Laboratory. Hospital Nacional de Parapéjicos (SESCAM), Finca La Peraleda s/n, E-45071 Toledo, Spain; Institut für Physiologische Chemie, Tierärztliche Fakultät, Ludwig-Maximilians-Universität, Veterinärstr. I3, D-80539 München,Germany

## Abstract

The mechanism underlying selective myelination of axons versus dendrites or neuronal somata relies on the expression of somatodendritic membrane myelination inhibitors (i.e. JAM2). However, axons still present long unmyelinated segments proposed to contribute to axonal plasticity and higher order brain functions. Why these segments remain unmyelinated is still an unresolved issue. The bifunctional lectin galectin-4 (Gal-4) organizes the transport of axon glycoproteins by binding to N-acetyllactosamine (LacNac) termini of N-glycans. We have shown that Gal-4 is sorted to segmental domains (G4Ds) along the axon surface, reminiscent of these long unmyelinated axon segments in cortical neurons. We report here that oligodendrocytes (OLGs) do not deposit myelin on Gal-4 covered surfaces or myelinate axonal G4Ds. In addition, Gal-4 interacts and co-localizes in G4Ds with contactin-1, a marker of non-myelinated nodes of Ranvier. Neither Gal-4 expression nor G4D dimensions are affected by myelin extracts or myelinating OLGs, but are reduced with neuron maturation. As in vitro, Gal-4 is consistently segregated from myelinated structures in the brain. Our data shape the novel concept that neurons establish and regulate axon membrane domains expressing Gal-4, the first inhibitor of myelination identified in axons, whose boundaries delineate myelination-incompetent axon segments along neuron development.

## Introduction

Axons are selectively myelinated while the somatodendritic membrane remains free of myelin insulation. Myelination does not require local stimulation or special geometric conditions, as surfaces of any shape can be myelinated in the absence of molecular inductors^1,2^. An explanation for this selective axon myelination has been recently reported, whereby the somatodendritic membrane expression of the Junction Adhesion Molecule 2 (JAM2) inhibits local myelination^3^. Even though most axons are selected for myelination, they are not completely covered with myelin. The formation and significance of long unmyelinated axonal tracts, which have been related to high axonal plasticity underlying complex cognitive functions^4^, remain open questions.

The extensive complexity of glycan determinants and their dynamic changes are linked to key functions of the nervous system^5–9^, including those related with myelin structure. Glycoconjugates as myelin-associated glycoprotein (MAG), a sialic acid-binding lectin (Siglec-4a)^10^, gangliosides GD1a/GT1b or integrins play essential roles in internode stabilization. An illustrative example is contactin which is transported to myelinated axon paranodes as its N-glycans remain immature. Upon N-glycan maturation to their form bearing LacNAc termini, contactin is sorted to non-myelinated nodes of Ranvier, where it interacts with other glycosylated components (i.e. neurofascin NF-186), and stabilizes the nodal structure ^11–13^.

Importantly, the detection of glycan endogenous receptors (lectins) in the nervous system has prompted the idea of a functional glycan/lectin pairing^14–16^. In particular, the reported expression of members of the galectin family, tissue lectins with β-galactoside binding capacity^17,18^, has stimulated the research of their functional implications in nervous system physiology. So far, galectin-1 (Gal-1)^19–23^, galectin-3 (Gal-3)^24,25^, and galectin-4 (Gal-4)^26,27^, have been related to axon growth and guidance, differentiation of microglia and oligodendrocytes, or nerve regeneration. LacNAc-dependent sorting of contactin suggested the participation of galectin interactions in myelin organization, according to our previous report of a directional Gal-4-based axon transport of LacNAc-bearing glycoproteins. In addition, Gal-4 is early sorted to the axon membrane, and it is expressed in discrete segments along axon surface of mature neurons^27^. The role of these membrane Gal-4-containing domains (G4Ds) is unknown, though their disposition in axons, reminiscent of nonmyelinated segments of cortical neurons^4,28^, and the fact that soluble Gal-4 retards oligodendrocyte maturation^26^, prompted our hypothesis that Gal-4 in G4Ds could produce localized myelination impairment of functional consequences.

Here we identify Gal-4 as the first inhibitor of local myelination in axons, and propose that its expression in axon membrane domains defines myelination-incompetent axon segments regulated along neuronal development.

## Results

### Gal-4 is sorted to axon membrane domains in mature neurons

Gal-4 in immature neurons is enriched at the membrane of the nascent axon^27^, and is secreted by non-classical mechanisms to the extracellular milieu, where it keeps oligodendrocyte precursors (OLPs) in their undifferentiated proliferative state, unable to synthesize myelin^26^. Even though hippocampal and cortical neurons reduce the expression and the secretion of Gal-4 when differentiated, the expressed lectin is sorted to discrete segments along the axonal plasma membrane, as indicated by Gal-4 immunostaining of non-permeabilized cultured neurons (Figure 1A, arrowheads). Similarly, exogenously induced HA-tagged Gal-4 (Gal-4-HA) is also transported and targeted to axon membrane segments (Figure 1B, arrowheads). Of note, when Gal-4-HA expression is induced in other cells of the nervous system that express very low levels of endogenous Gal-4, such as astrocytes, non-permeabilized immunostaining for HA is observed as membrane “patches” (Figure 1C, arrowheads). Such a consistent mechanism for the formation and maintenance of Gal-4- containing membrane domains (G4Ds) in axons suggested a potentially important role in neuronal physiology.

**Figure 1.**
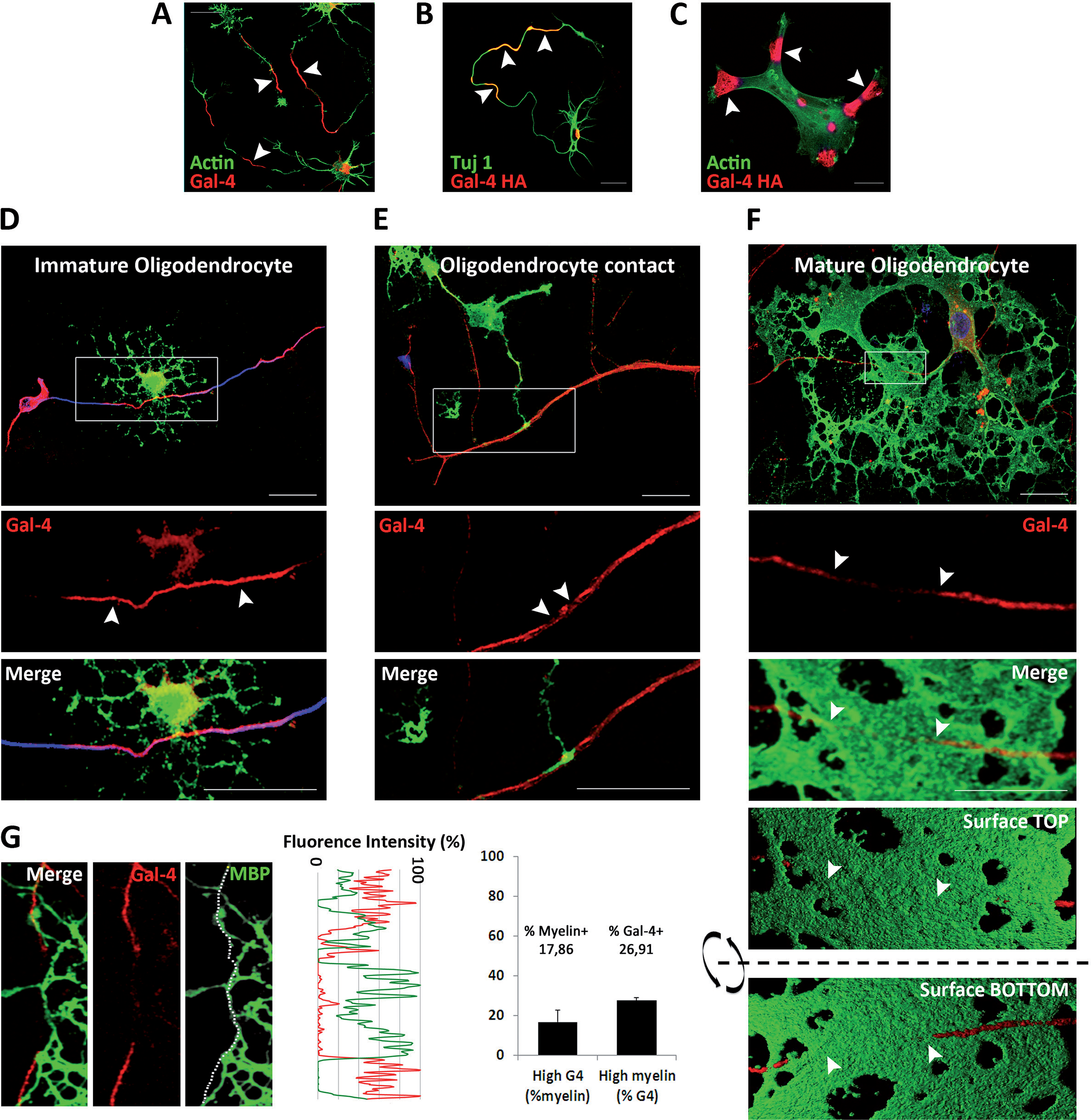
Axonal G4Ds are not myelinated in neuron/oligodendrocyte co-cultures. **(A)** Hippocampal neurons (72 hiv) were immunolabelled for Gal-4 (red) without permeabilization. Gal-4 is detected in discrete segments (arrowheads) along the axonal plasma membrane. **(B)** HA-tagged Gal-4 (Gal-4-HA) transfected hippocampal neurons were immunolabelled for HA (red) without permeabilization. Gal-4-HA is also detected in axon membrane segments (arrowheads). **(C)** Gal-4-HA transfected astrocytes were immunolabelled for HA (red) without permeabilization. Exogenously expressed Gal-4-HA forms “patches” on the cell surface (arrows), reinforcing the concept of functional Gal-4- containing membrane domains. **(D-G)** Hippocampal neurons and oligodendrocytes were cocultured for different time periods. The ICC for Gal-4 (red) was performed under non-permeabilizing conditions to detect Gal-4 associated to axon surface. Cells were then permeabilized and immunolabelled for acetylated tubulin (blue) and MBP (green) to label axons and myelin, respectively. **(D)** The amount of Gal-4 associated to the axon membrane is not affected by the contact with oligodendrocytes at early stage of differentiation (arrowheads). **(E)** Axon points where initial contacts of myelinating oligodendrocytes occur are devoid of membrane Gal-4 (arrowheads). **(F)** At more advanced stages of myelination in vitro, membrane Gal-4 is absent in myelinated axon segments (arrowheads). A 3D reconstruction from a stack of Z-planes processed to simulate opaque surfaces show that the myelin sheet (green) entirely wraps the portion of the axon devoid of Gal-4 (red) as seen from the top and bottom surfaces of the reconstructed image. **(G)** Quantification of axon membrane Gal-4 and MBP, measured as percentage of the maximum fluorescence intensity in each channel. Axon segments in which Gal-4 levels are above 50% present significantly low levels of MBP, and viceversa (values are means + standard deviation (SD) of three experiments; n=49). (Scale bars 25 μm).

### Myelinated axon segments express very low Gal-4 membrane levels

Considering that secreted Gal-4 retards OLG maturation and myelin formation^26^, the segmented disposition of G4Ds suggested that Gal-4 could impede local myelination of those domains. To approach this question, we studied G4Ds in myelinating co-cultures of hippocampal neurons and OLGs, using myelin basic protein (MBP) labelling to assess OLG maturation. G4Ds were observed on axon membranes independently of eventual direct contact with non-myelinating OLGs. Figure 1D shows a representative image, evidencing that the portion of an axon growing on top of an immature OLG expresses strong membrane labelling for Gal-4 (Figure 1D, arrowheads). In contrast, the initial contact points between differentiated OLG extensions and axons are mostly devoid of Gal-4 (Figure 1E, arrowheads). As co-cultures proceeded in time, longer axon segments appeared myelinated by fully mature OLGs, coinciding with the absence of axon membrane Gal-4 (Figure 1F, arrowheads). After transforming confocal signal to show opaque surfaces (see methods for details), views from the top (Figure 1F, Surface Top) and from the bottom (Figure 1F, Surface Bottom) of the culture corroborate that the axon portion fully enwrapped by myelin (delimited by arrowheads) is practically devoid of Gal-4. Quantitatively, axon segments expressing high Gal-4 content, present in average 17,9 % of the maximum MBP (myelin) labelling (Figure 1G). In correlation, high myelin axon segments present in average 26,9% of the maximum Gal-4 label (“high content” of Gal-4 or MBP is considered when their respective labels are higher than 50% of each maximum fluorescence intensities, standardized to 100%).

### Surface-bound Gal-4 repels myelin deposition

Co-culture experiments corroborated that G4Ds are essentially non-myelinated. However, they do not clarify whether it is Gal-4 itself, or any other component of those domains, that drive(s) myelin distribution along the axon. In order to test the direct contribution of bound Gal-4 to myelin distribution, we cultured OLGs on glass coverslips patterned with parallel alternated stripes, covered or not, with recombinant galectin. This display challenges OLGs to select the preferred surface to myelinate, resembling the situation they would find in cocultures and likely *in vivo.* FITC was used together with galectins or alone (control) to visualize the stripes (bright stripes). MBP-labelled areas in galectin-containing and galectin-free stripes were measured and compared as percentages of the total. After 48 hours in culture, mature OLGs cultured on Gal-4-parallel stripes presented an irregular shape, suggesting a repulsive effect to myelin deposition, which is preferentially spread on Gal-4-free surface (Figure 2B, black stripes). In fact, 68.9% of myelin was formed on Gal-4-free stripes opposed to Gal-4-covered stripes (Figure 2F, Gal-4), while 51.2% did when opposed to control FITC-covered stripes (Figure 2A and 2F, FITC). Similar results to those of Gal-4 were obtained if stripes were covered with recombinant N-terminal domain of Gal-4 (Gal-4N, Figure 2C). In this case, 64.6% of the myelin was formed on Gal-4N-free stripes (Figure 2F, Gal-4N). In contrast, stripes covered with the recombinant C-terminal domain of Gal-4 (Gal-4C, Figure 2D) did not trigger any effect, as 45.2% of the myelin was formed on Gal-4C-free stripes (Figure 2F, Gal-4C). The specificity of the Gal-4 effect on myelin formation was further revealed by the lack of effect of different full-length galectins, as shown for recombinant Gal-3 (46.3% of myelin formed on galectin-free stripes; Figures 2E and 2F, Gal-3).

**Figure 2.**
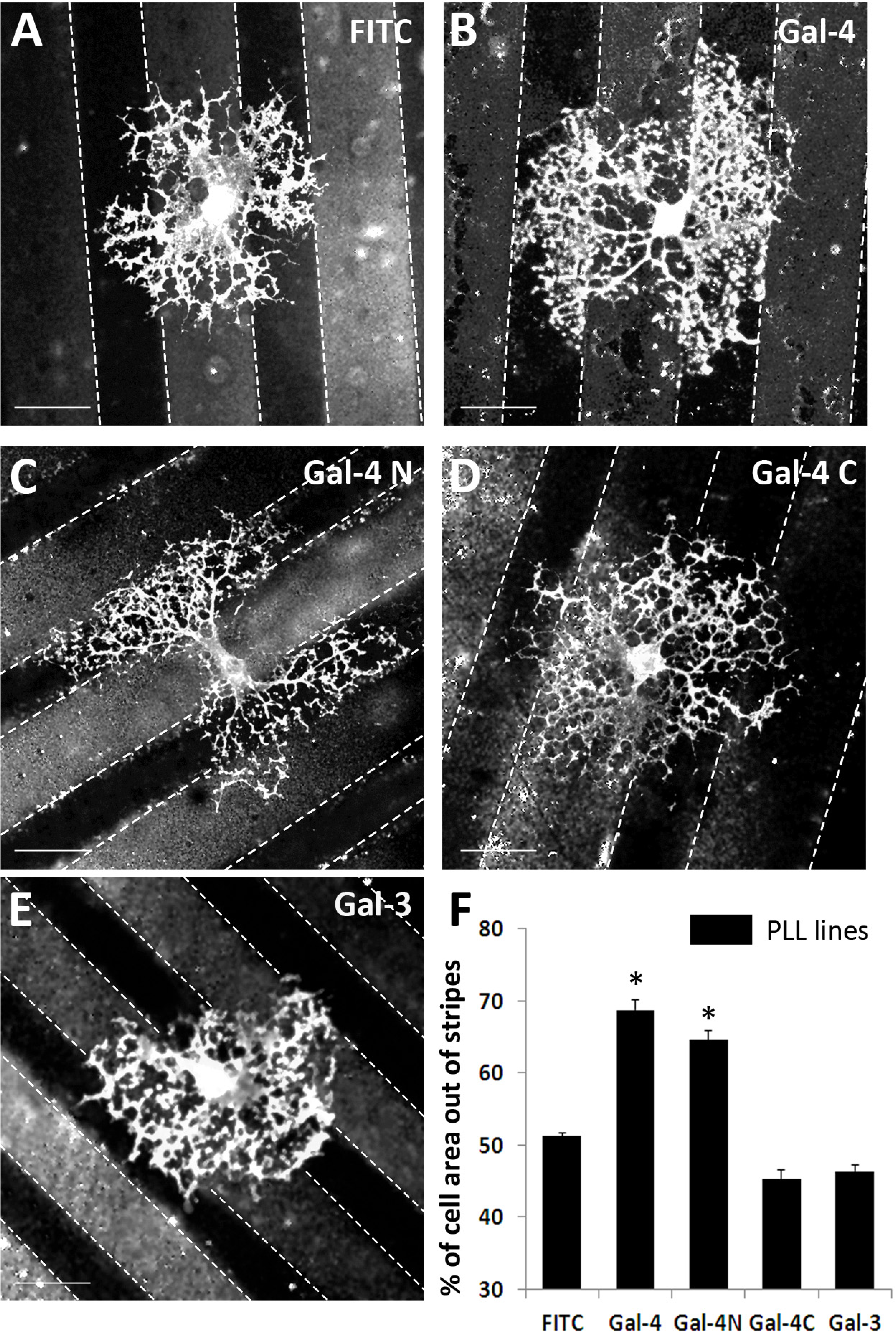
Gal-4 covered substrates reject myelin deposition. **(A-E)** OLGs were allowed differentiate for 72 hours on coverslips with parallel stripes covered with recombinant Gal-4, Gal-4N, Gal-4C and Gal-3. FITC (brighter stripes) was used together with galectins to evidence galectin-covered stripes **(B-E),** or alone as negative control **(A).** Cells were immunolabeled for MBP to visualize myelin. OLGs preferentially grow their myelinating processes on Gal-4-free **(B)** and Gal-4N-free **(C)** substrates, while Gal-4C does not produce any effect **(D)**. The presence of Gal-3, used as control, does not affect myelin deposition **(E)**. Quantitative analysis of the repulsive effect exerted by Gal-4 on mature OLGs is shown in **(F)**. The bar graph shows the percentage of MBP-positive area on stripes covered with PLL alone, that is, out of galectin-containing stripes. Values are mean + SD of three experiments (20 cells/experiment) (*p<0.005, Student’s /-test); (Scale bars 25 μm).

These results indicate that bound Gal-4 is capable to preclude myelin deposition by interaction with myelinating OLGs, and that this repulsive effect likely resides in Gal-4’s N-terminal domain. Although it cannot be ruled out that other G4D components could play a role in myelin organization along the axon, our data support that Gal-4 is a direct modulator of that process.

The identification of G4Ds as non-myelinated axon domains suggested that they could concentrate glycoproteins normally found in non-myelinated nodes of Ranvier between myelinated internodes. Both in cultured hippocampal (Figure 3A) and cortical (not shown) neurons, nodal GPI-anchored glycoprotein contactin (Figure 3A, arrowheads) extensively localize in G4Ds. Furthermore, using extracts of PC12 cell line that express both proteins, we show that anti-contactin antibodies co-immunoprecipitate Gal-4 (Figure 3B, IP contactin, WB Gal-4). Fittingly, the reverse experiment showed that Gal-4 antibodies co-immunoprecipitated contactin (Figure 3B, IP Gal-4, WB contactin).

**Figure 3.**
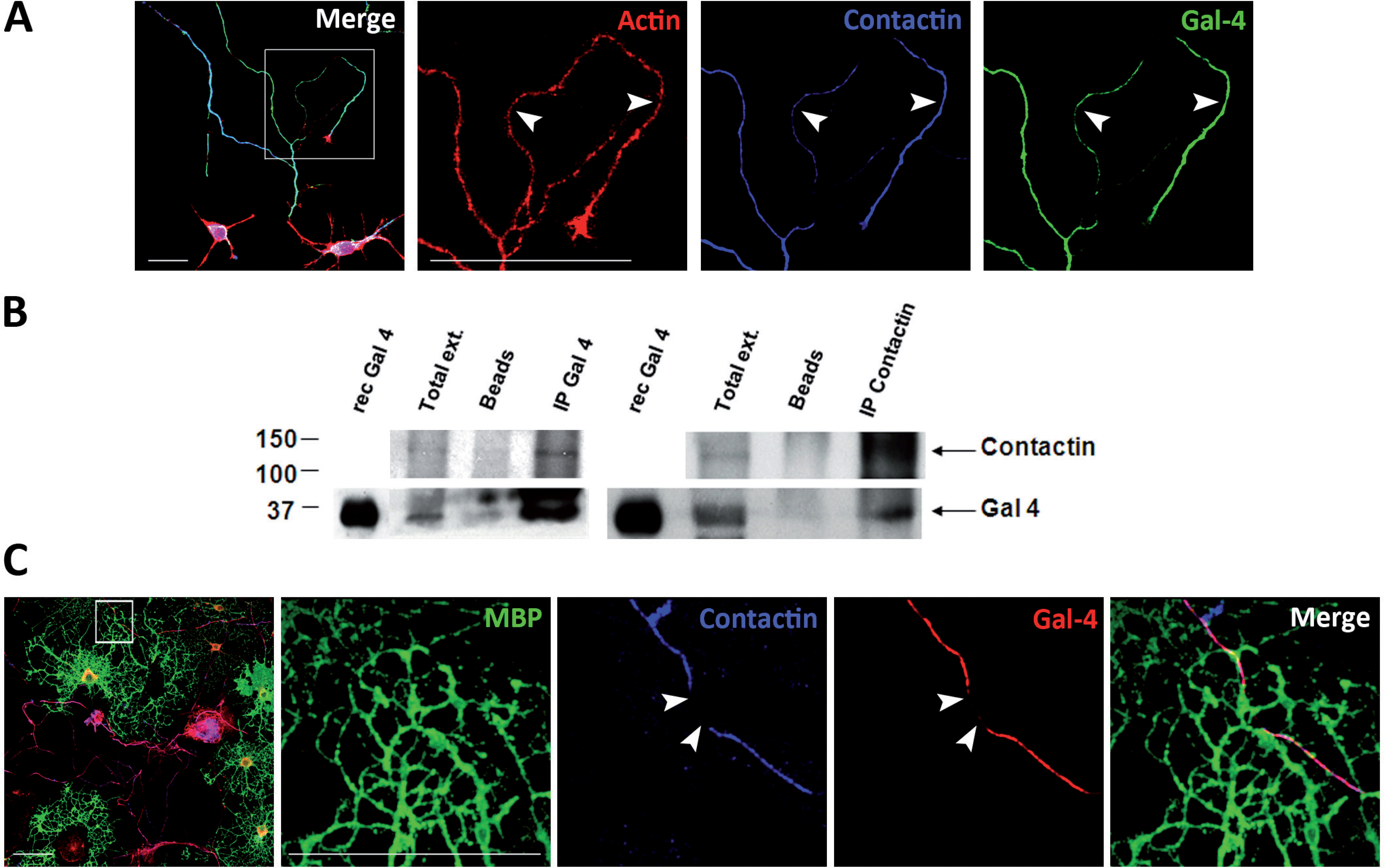
Nodal marker contactin interacts and co-localizes with Gal-4 in nonmyelinated axon segments. **(A)** Hippocampal neurons (72 hiv) were immunolabelled for Gal-4 (green) and contactin (blue) without permeabilization. Gal-4 and contactin colocalize in discrete segments (arrowheads) along the axonal plasma membrane. F-actin (red) was labelled with phalloidin to visualize neurons. **(B)** PC12 cell cultures were reversibly crosslinked, lysed, and protein extracts (Total ext.) were used to immunoprecipitate Gal-4 or contactin. The immunoprecipitated complexes were separated by SDS-PAGE and detected by WB. Recombinant Gal-4 (rec Gal-4), and incubations with protein-A or –G Sepharose beads in the absence of antibodies (Beads) were used as controls. Anti-Gal-4 precipitated contactin (IP-Gal-4, WB Contactin). Consistently, anti-contactin 1 precipitated Gal-4 (IP Contactin, WB Gal-4), demonstrating the interaction of both proteins. **(C)** Hippocampal neurons and oligodendrocytes were co-cultured, fixed and analyzed by immunofluorescence. Labellings for Gal-4 (red) and contactin (blue) on axon membrane were performed under non-permeabilizing conditions. Cells were then permeabilized and immunolabelled for MBP (green) to visualize myelin. Myelinated axon segments present low levels of Gal-4 and contactin (arrowheads) (Scale bars 25 μm).

Gal-4/contactin co-localizations were observed in cultures of purified neurons. So it was still possible that such interactions would be altered by myelination. Immunocytochemical labelling of neuron/OLG co-cultures revealed that Gal-4 and contactin (Figure 3C), colocalize in non-myelinated axon tracts. Those tracts in close contact with myelin, labelled with antibodies anti-MBP, were mostly devoid of Gal-4, and contactin (Figure 3C, arrowheads). Gal-4 interaction with the nodal marker contactin, and its absence in myelinated segments, further support the notion that G4Ds are excluded from myelination and contribute to organize the discontinuous structure of myelin along the axons.

### Myelin does not affect G4Ds membrane display

Myelination is a dynamic process, thus final myelin arrangement would require the variation of G4Ds dimensions and disposition. We hypothesized that the contact with myelin, as it expands laterally, could reduce G4Ds by inducing the elimination of Gal-4 from the axon membrane. To test this concept, 48 hiv hippocampal neurons were treated for different time periods with freshly prepared myelin extracts, and the levels of membrane Gal-4 were measured as average fluorescence intensity in immunolabelled cultures. Independently to the duration of the treatment or extract concentration (up to 24 hours and 12 μg total protein shown in Figure 4A), no difference in Gal-4 membrane labelling was detected between treated and untreated cultures (Figure 4A, bar graph). To further corroborate these results, identical treatments were applied to astrocytes induced to express Gal-4-HA (Figure 4B). Similarly to neurons, Gal-4-HA membrane patches appeared unaffected by the treatment with myelin extracts, as fluorescence intensity quantification showed non-significant differences with control cultures (Figure 4B, bar graph). Even though the average Gal-4 content of the membrane was not affected by myelin extracts, it was still possible that its distribution could be altered. To check this, G4Ds lengths of myelin-treated neurons were measured and compared with controls. Myelin treatment did not change significantly the dimensions of G4Ds. In detail, average lengths in myelin-treated and non-treated cultures were 28.9 ± 0.84 μm and 28.4 ± 1,28 μm, respectively (Figure 4C, upper graph). Nevertheless these results did not preclude the possibility of G4Ds modulation by myelin presented on OLG surface, in contrast to soluble extracts. To address this question, a parallel quantification of G4D length was performed in 21 div neurons co-cultured with myelinating OLGs for the last 7 div, and compared to that of control 21 div neurons. As for myelin extracts, no differences in G4D length due to myelinating OLGs could be detected (Figure 4C, lower graph).

**Figure 4.**
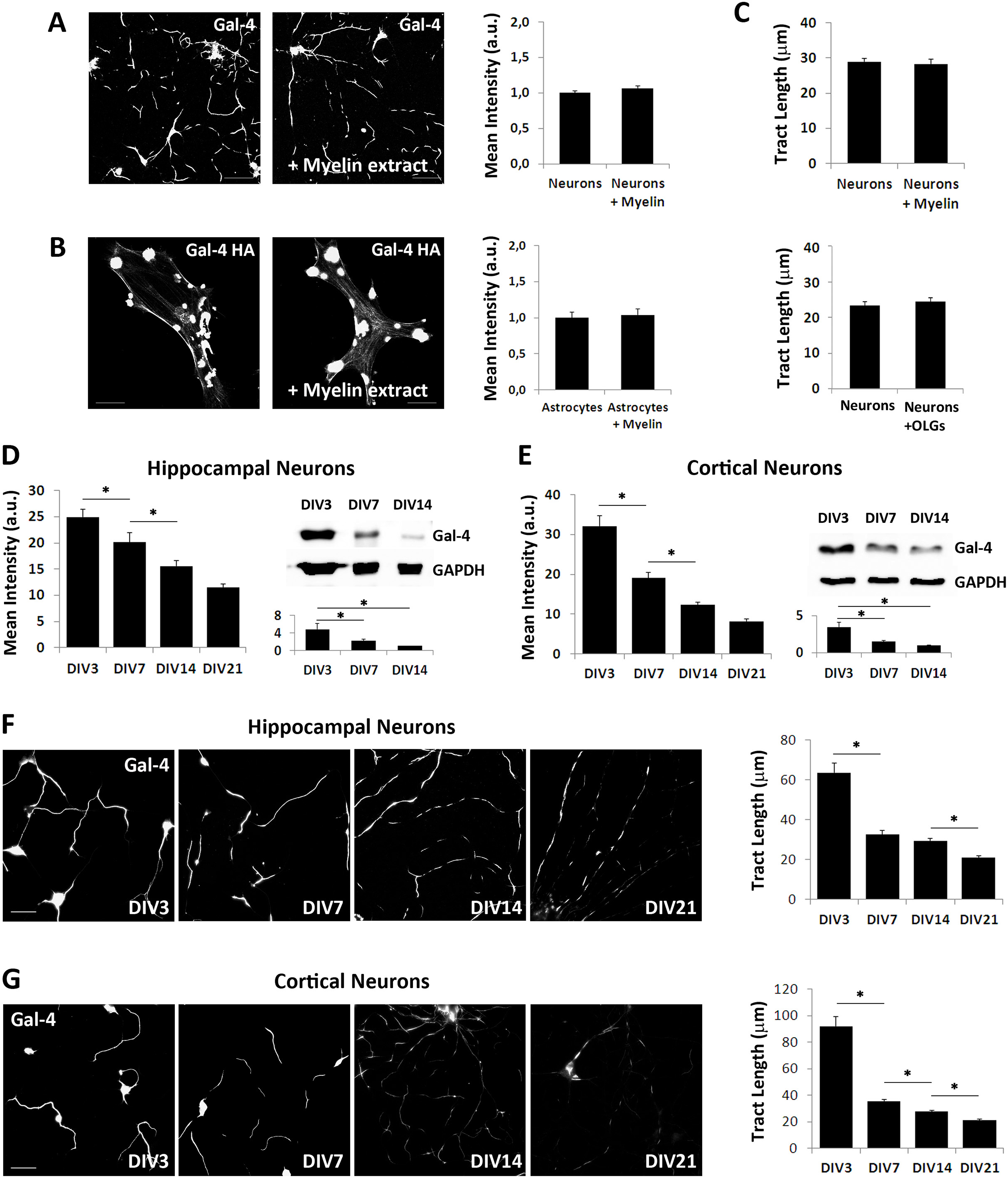
Gal-4 expression and G4D dimensions are regulated along neuronal maturation independently of myelin. **(A)** Hippocampal neurons (48 hiv) and **(B)** Gal-4-HA-expressing astrocytes, treated for 24 hours with myelin extracts (+Myelin extract) or untreated (left panels in A and B), were immunolabelled without permeabilization for Gal-4 **(A)** and HA **(B)**. The levels of membrane Gal-4 and Gal-4-HA were measured as average fluorescence intensity for neurons (bar graph in A) and astrocytes (bar graph in B), respectively. (C) G4Ds lengths of myelin-treated neurons were measured and compared with controls. No significant differences were observed for any of the measured parameters. **(D-E)** Hippocampal and cortical neurons were immunolabelled without permeabilization for Gal-4, and the average fluorescence intensity was measured (left bar graphs in D and E for hippocampal and cortical cultures, respectively). From 3 to 21 div, membrane Gal-4 mean fluorescence intensity significantly decreases. Upper right panels show representative WB for total Gal-4 expression in extracts of hippocampal **(D)** and cortical **(E)** neurons at different times in culture. GADPH was used as loading control. Lower right bar graphs in **(D)** and **(E)** display average Gal-4 densitometric measures referred to the minimum values at div 14, arbitrarily considered as 1 in each case. Experiments were performed in triplicate. A significant reduction of total Gal-4 expression is observed between 3 and 14 div for both neuron types. Hippocampal **(F)** and cortical **(G)** neuron cultures were immunolabelled without permeabilization for Gal-4. The lengths of Gal-4 positive axon segments were measured in each condition. The bar graphs show acute drops in G4D length at 7 div, and slow decreases later on (n=2951 for hippocampal neurons and n=2148 for cortical neurons) (*p<0.05, Student’s t-test). Experiments were replicated four times for each cell type. All bar graphs display mean values and standard errors. (Scale bars 25 μm).

### Neurons modulate Gal-4 expression and G4D size during axon development independently of myelin

Using Gal-4 display as read-out parameter, our results show that the contact with myelin does not modulate G4Ds on the axonal membrane, pointing to a neuronal control of those domains that could be modulated during axon development. Such a regulation would imply modifications of Gal-4/G4Ds with time in the absence of myelin or OLGs. To prove this we first followed Gal-4 membrane expression in hippocampal and cortical neuron cultures with time. From 3 to 21 div, membrane Gal-4 mean fluorescence intensity significantly decreases in the range of 50% and 70% for hippocampal and cortical neurons, respectively. Hippocampal neurons show a linear descending trend showing constant decreases at 7, 14 and 21 div, while cortical neurons show a pronounced drop between 3 and 7 div (around 59%), and slower decreases at 14 and 21 div (Figs. 4D and 4E, bar graphs on the left). These results correlate with the reduction trends of total Gal-4 with time in culture measured by Western blot (WB) for both, hippocampal and cortical neurons (Figures 4D and 4E, blots and bar graphs on the right). According to these data, the decrease of membrane Gal-4 with neuron maturation is due, at least in part, to a reduction of its expression rate, although these changes in Gal-4 expression do not necessarily imply modifications in G4Ds distribution or size. To check this we measured G4D length evolution with time in hippocampal and cortical neuron cultures (Figures 4F and 4G). At 3 div G4D length is maximum, being longer in cortical (95.5 ± 6.8 μm) than in hippocampal (61.36 ± 4.1 μm) neurons, and covering in some cases most of the axon length. At 7 div an acute drop in G4D length occurs, reaching averages that range 30 μm in both neuron types. These values are maintained up to 14 div, to undergo a further significant decrease to approximately 20 μm at 21 div (Figure 4F and 4G, bar graphs). These results demonstrate that neurons can modulate axon membrane G4Ds along their development, independently of any interaction with OLGs or myelin.

### Gal-4 and MBP (myelin) expression are segregated in rat brain cortex

According to our results *in vitro,* we hypothesized a negative correlation of Gal-4 and myelin expression in brain tissue. To corroborate this, IHC labellings for Gal-4 and MBP were performed on vibratome slices of P3 and P30 rat brains (Figure 5A). Previous reports^26^ showed by WB that Gal-4 expression is high in P3 rat brains and decreases at P30 or later. The opposite was shown for MBP, whose expression is very low in P3 brain, while P30 brains express moderate quantities of the protein. In agreement with these results, MBP labelling in P3 brain was fairly undetectable, while a descending gradient of MBP labelling was evident in P30 rat brains from the *striatum* to the cortex surface (Figures 5B, 5C, red channel; 5D, black line). In contrast, Gal-4 is expressed in neurons of the cortical plate (CP) and layers IV/V of P3 brains (Figure 5B, green channel), labelling the somata and fibres projecting to the brain surface (Figure 5B, insets 1-2 arrowheads). In P30 brains Gal-4 labelling augments from deeper layer to the brain surface, and defines an inverted expression profile with respect to that of myelin (Figure 5D, gray line). Gal-4 levels at myelinated *striatum* and cortical layer VI are very low and restricted to somata (Figure 5C, inset 4, green channel). At layers IV and V, Gal-4 labelling is higher and localizes on neuronal somata and ascending fibres (Figure 5C, insets 2-3, arrows and arrowheads, respectively), while at the outermost, scarcely myelinated layers I-III, Gal-4 expression is mostly confined to axon fibres (Figure 5C, red channel; inset 1, arrowheads).

**Figure 5.**
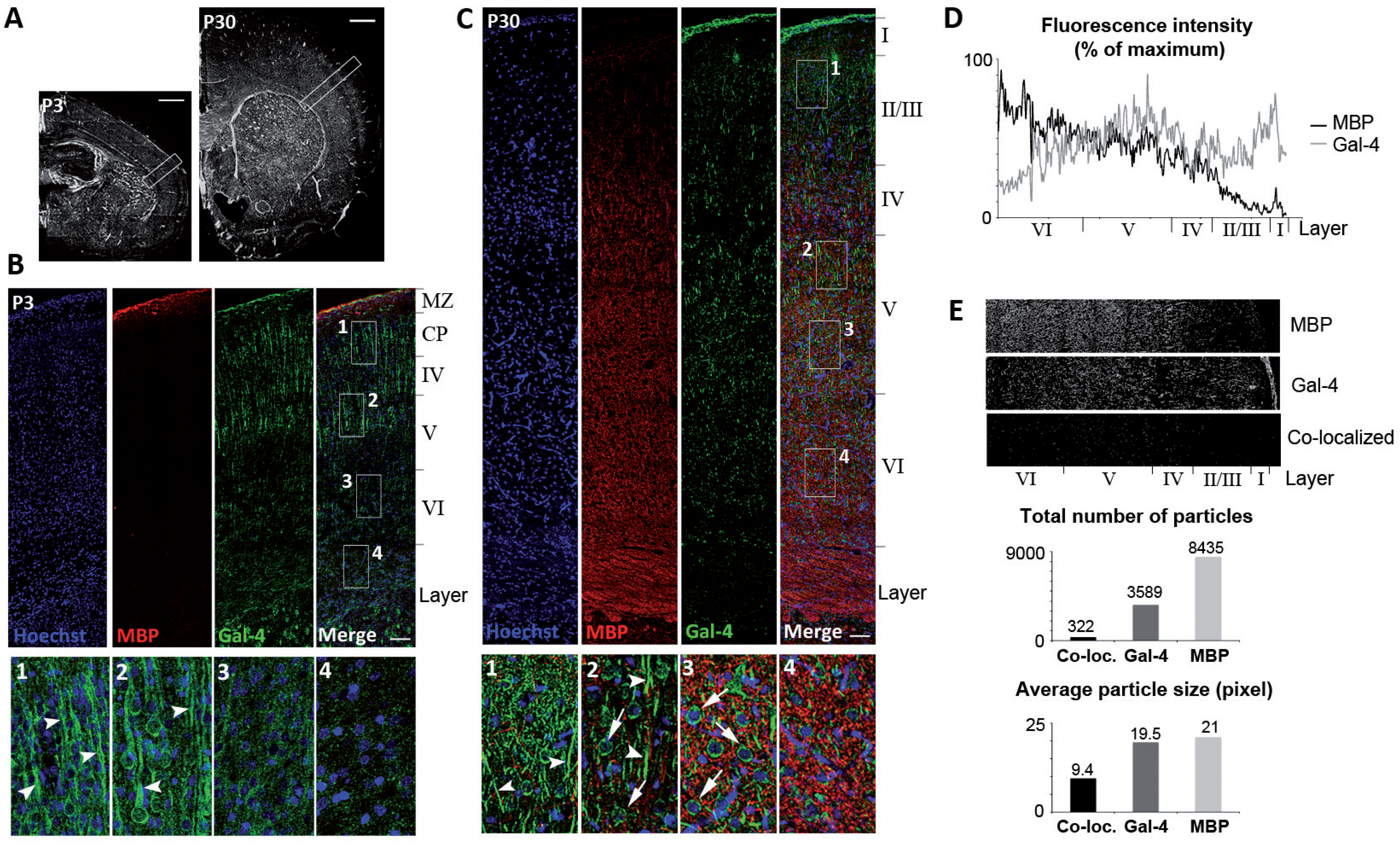
Gal-4 distribution along the brain cortex inversely correlates with myelin. **(A)** Low-magnification photomicrograph of a P3 (left) and P30 (right) rat cerebral hemisphere stained with Hoechst. White boxes indicate the regions within the somatosensory cortex shown in B and C. (Scale bars 500 μm). **(B-C)** Maximum intensity projections of confocal mosaic stacks obtained from a P3 **(B)** and P30 **(C)** rat cortex. The sections were immunolabeled for MBP (red) and Gal-4 (green). Nuclei were stained with Hoechst (blue). Numbered boxes (1-4) representing different cortical depths are shown at higher magnification in the bottom images. Note the greater expression of Gal-4 in the upper layers (boxes 1 and 2) compared to the bottom layers (boxes 3 and 4), both at P3 and P30. Gal-4 is present in fibers (arrowheads) as well as cell bodies in the upper layers, while its expression is largely confined to cell bodies at bottom layers (boxes 3 and 4, arrows in C). (Scale bars 100 μm). **(D)** MBP and Gal-4 fluorescence intensity (for the P30 cortex shown in C) was measured from the pia to the white matter, its profile was plotted and the values normalized to the maximal intensity obtained for each channel. Note how in deeper cortical layers (VI) the intensity for MBP is maximal while Gal-4 displays its lowest levels. On the contrary, the upper scarcely myelinated layers (I-III) show abundant Gal-4 fluorescent intensity. **(E)** MBP and Gal-4 positive particles (from the P30 cortex shown in C) shown in the upper and central images, respectively, were analyzed to determine their degree of colocalization. Co-localizing particles are shown in the lower image, and the quantification of their number and their average size is shown in the bar graphs below, indicating that colocalization level is low, and that the average size of co-localizing particles is significantly smaller compared to Gal-4 or MBP particle.

Considering the inverted expression profiles of Gal-4 and MBP (myelin) in brain cortex, we further explored the degree of co-localization of both molecules. Thresholded labels of Gal-4 and MBP channels were used to create maps of positive particles (see methods for details) as the ones shown for red and green channels of figure 5C (Figure 5E, upper and central images). The combination of those maps showed that co-localizing particles were less than 10% of Gal-4 particles (Figure 5E, lower image and upper graph), and that their average size was more than 50% smaller than that of Gal-4 or MBP particles (Figure 5E, lower graph). Of note, those quantifications were made using maximum projection of many confocal z-planes, thus part of those co-localized particles could be due to optical superposition rather than to actual co-localization. In fact, similar studies using single confocal z-planes of different cortical layer fields showed that co-localizing particles were much fewer (<3%) and smaller (<26%) in proportion to Gal-4 or MBP particles (Supplementary Figure S1).

In all, brain IHC studies confirm our results from *in vitro* experiments presented so far, and support the concept that Gal-4 expression on neuronal membrane locally excludes the formation of myelin.

## Discussion

OLGs require neither specific geometric conditions^1,2^, nor inductive substrates to form myelin, indicating that selective myelination of axons is not due to the presence of local myelination inductors. On the contrary, recent studies have shown that the somatodendritic compartment of neurons is not myelinated due to the expression on the membrane of a myelination inhibitor JAM2^3^ which would explain why axons are selected for myelination. Nevertheless, axons are not completely myelinated, and the formation and function of long non-myelinated discontinuities along the axon are still unresolved issues.

We reported that mature neurons express Gal-4, a galectin that is specifically sorted to axon membrane segments in a sulfatide-dependent manner^27^. As documented here, exogenous HA-tagged Gal-4 expressed in neurons (Figure 1B) displays the same pattern on the axon membrane than the endogenous Gal-4 (Figure 1A), or is sorted to membrane “patches”, when expressed in astrocytes that express very low levels of endogenous Gal-4 (Figure 1C). This precludes the possibility of technical artifacts, and reinforces the concept of functional Gal-4-containing axon membrane domains.

Taking into account that Gal-4 secreted by immature neurons retards myelination by blocking the maturation of OLPs^26^, and that G4D display along the axon are reminiscent of the non-myelinated axon segments of the outer neocortical layers^4^, we hypothesized that Gal-4 could locally impair myelination of G4Ds, contributing to the formation of nonmyelinated axon tracts. In fact, myelinated portions of the axon are practically devoid of G4Ds, both at initial and at advanced myelination stages (Figure 1E and 1F, Supplementary video), while these domains are not excluded from axon contacts with pre-myelinating OLPs (Figure 1D). Whether Gal-4 was directly responsible for G4Ds exclusion from myelinated segments remained obscure. Unfortunately, knocking-down Gal-4 by interference RNA was not appropriate to approach this issue because low level of Gal-4 significantly retards axon growth, as we previously shown^27^. Alternativelly, cell culture surfaces with parallel microstripes, covered or not with recombinant Gal-4, were suited to mimic the scenario in which OLGs are challenged to myelinate G4Ds or Gal-4-free axon segments. Under such conditions, OLGs preferentially myelinated Gal-4-free stripes (Figures 2B and 2F).

Moreover, recombinant N-terminal domain of Gal-4 was sufficient to preclude myelin deposition (Figure 2C and 2F). In consequence, it appears that Gal-4 presence is sufficient to keep myelination excluded from G4Ds, conferring functional significance to these domains.

The bivalent Gal-4 is capable to bridge glycans in a rather stable manner, stabilizing/organizing “superrafts” and yielding vesicle aggregation, proof for the concept of higher-order microdomain formation^29,30^. We show here that Gal-4 within G4Ds co-localizes and interacts with contactin (Figure 3), whose N-glycans need to acquire complex LacNAc termini for a correct targeting to nodes of Ranvier^6, 11–13^. The fact that Gal-4 has strong binding affinity for LacNAc presented in clusters^31–34^, could underlie contactin recruitment to myelin-excluding G4Ds.

The idea that MBP labelling (myelin) is absent on Gal-4-expressing neuron membrane, and that this is maintained at different developmental stages, is strongly supported by comparison of the expression profiles of both molecules in brain tissue. At P3, when myelination is beginning, Gal-4 is present in neurons of the CP and developing layers IV and V (upper part), but not in layer VI or striatum, two zones that will become heavily myelinated later on (Figure 5B). Consistently, when myelination is quite advanced, as shown in P30 brains, barely myelinated layers II-III, and layers IV-V maintain different patterns of Gal-4 expression, while myelinated layer VI and striatum show very low expression of the galectin (Figure 5C). Those inverted Gal-4 and MBP expression profiles along the axis from white matter to brain surface (Figure 5D), are consistent with the almost complete segregation of Gal-4- and MBP-labelled sub-cellular structures, evidenced by colocalization analyses (Figure 5E; Supplementary Figure S1). The analysis of brain tissue thus reinforces the conclusion obtained *in vitro* that Gal-4 excludes myelin deposition, pointing to a functional role in the developmental organization of myelin.

Axon myelination proceeds by lateral expansion of the concentric myelin sheaths until myelin configuration is established, with myelinated segments (internodes) separated by nodes of Ranvier or by the recently reported long non-myelinated tracts in cortical layers I-III ^4,28^. In either way, final myelin arrangement would require shrinkage of G4Ds and/or changes in their disposition along the axon. We first thought that myelinating OLGs could modulate this process, and proposed that the contact of myelin with one or more G4D components would induce the elimination of Gal-4 (and probably other molecules) from the axon surface through the activation of local endo/exocytosis. However, no changes in G4D quantity or dimensions were observed in axons exposed to myelin extracts (Figure 4A, and 4C upper graph). Importantly, G4Ds were neither affected by myelinating OLGs (Figure 4C, lower graph), which precludes the possibility that G4D modulation could depend on myelin presentation on the OLG surface. In all these results pointed to a myelin-independent neuronal regulation of these domains with neuron differentiation. In support of this hypothesis, total Gal-4 expression in hippocampal and cortical neurons significantly decreases with time in culture, which is reflected by a concomitant reduction of the amount of galectin detected on neuronal membranes (Figures 4D and 4E). In addition, G4Ds become shorter as neurons mature in the absence of any contact with myelin (Figures 4F and 4G), indicating that the arrangement of these domains, which influences myelin deposition, is regulated by neurons. It is important to note that the highest values of Gal-4 expression and of G4Ds average length in cultured neurons correspond to the fast axon growth period (up to 3 div), when low myelination must be maintained in order to allow the efficient elongation of the axons. Within the next five div, Gal-4 expression undergoes a severe reduction and, from that time point on, G4Ds continue a further slow-rate shortening, in coincidence with the time period when myelination actively progresses.

Furthermore, spontaneous synaptic activity increases with neuron differentiation in a time pattern similar to that observed for G4D variation. In cultures of dissociated neocortical neurons the number of spiking neurons augment significantly between 6 and 15 div^35^. That neuronal activity regulates CNS myelination is widely accepted^36–38^, and the local signalling mediated by axon synaptic vesicle release effect on OLGs, has been proposed as an important mechanism for this regulation^39-41^. Considering the contemporary variations of G4Ds and spontaneous synaptic activity in cultured neurons, it is appealing to speculate that, independently of OLGs, changes in spontaneous synaptic release could drive the modifications of Gal-4 and G4Ds that organize axon membrane for myelination. This would be compatible with the activation of OLGs by synaptic vesicle release mentioned above. The presented results and arising reasoning give further research into this question a clear direction.

## Methods

### Antibodies

The following antibodies were used: acetylated α-tubulin and βIII-tubulin (Tuj1, Sigma), MBP (Abcam), HA (Roche), contactin 1 (SCBT), and Gal-4 and -3 (in-house production). These two anti-galectin antibodies were affinity-purified to exclude cross-reactivity using the respective galectin as bead-immobilized ligand^42^. F-actin was detected with phalloidin (Molecular Probes). Appropriate anti-species secondary antibodies were from Molecular Probes.

### Recombinant galectins

Human galectin-4 (Gal-4) and its two separate carbohydrate recognition domains corresponding to a.a.1-150 and 176-323, referred to as N- and C-terminal domains (Gal-4N/C), respectively, were obtained by recombinant production, purified by affinity chromatography on lactosylated Sepharose 4B, and controlled for purity by 1D- and 2D-gel electrophoresis and mass spectrometry, and for activity by solid-phase and cell binding assays^31,43,44^.

### Cell cultures

Rat hippocampal and cortical neurons, primary astrocytes and PC12 cells were cultured as described previously^27^. Primary OLG cultures were prepared from the neocortex of 1-3-day-old rats, as previously described^45^. The purified OLGs were maintained in serum-free growth medium consisting of DMEM/Ham’s F12 (Gibco) (1:1) containing sodium selenite (30 nmol/l), putrescine (1 mg/l), insulin (5 mg/l), hydrocortisone (20 nmol/l), progesterone (20 nmol/l), transferrin (25 mg/l), biotin (10 nmol/l), 3,3’,5-triiodo-L-tyronine (30 nmol/l), bovine serum albumin (BSA 1mg/ml), supplemented with PDGFAA (5 ng/ml) and bFGF (10 ng/ml) for 48 h. Cells were then allowed to mature by growth factor withdrawal.

Neuron/oligodendrocyte co-cultures were performed as previously described by Gardner and collaborators ^46^ with some modifications. Oligodendrocyte were seeded onto 7 div cultured neurons, and cultured in MEM-HS and serum-free OLG growth medium in a 3:1 ratio. OLG medium proportion was gradually increased every 3 days by adding 1ml each time. After 17 days of co-culture, cells were fixed and analysed by ICC.

### Neuron-free myelination assay on galectin-stripped carpets

Substrate-challenge assays were set up by coating the surface of glass coverslips with recombinant Gal-4, Gal-4N, Gal-4C, and Gal-3 in parallel stripes. Silicon matrices with parallel micro-channels were used as described previously^47^. Briefly, solutions containing the Gal-4 proteins (1 mg/ml) and FITC (1 μg/ml; Sigma-Aldrich) were injected into the matrices and incubated for three hours at room temperature. Striped substrata covered with FITC alone were used as controls. Oligodendrocyte precursors were seeded and cultured for 12 hours at 37 °C in proliferation medium, and cultured for further 48 hours after growth factor withdrawal. Cells were then fixed and analyzed by ICC. Myelin total area and myelin area on galectin-free substrates (PLL lines) were measured using ImageJ software (NIH).

### Immunocytochemistry

For ICC analysis, cells were fixed with 4% paraformaldehyde for 15 minutes at RT, aldehyde groups were quenched with 50 mM ammonium chloride for 5 minutes. When indicated, membranes were permeabilized with 0.1% Triton X-100 in PBS during 5 minutes. Cells were incubated in blocking solution (2% FBS, 2% BSA, 0.2% gelatin in PBS) for one hour, followed by incubation with primary antibodies for one hour at RT or overnight at 4°C. After extensive washing, cells were incubated with appropriate secondary antibodies for one hour. For simultaneous detection of neuronal membrane-associated Gal-4, MBP and/or acetylated α-tubulin, a first ICC for Gal-4 detection was performed under non-permeabilizing conditions. Cells were then fixed again, permeabilized, and a second ICC was performed to detect MBP and/or acetylated α-tubulin. Coverslips were then mounted on glass slides with Mowiol (Calbiochem) using DABCO (Sigma) as anti-fade agent. Images were acquired on a confocal microscope (Leica SP5) or/and an epifluorescence microscope (Leica DM5000B).

### Cell extracts and Western blotting

Total cell extracts for biochemical analysis were obtained from hippocampal and cortical neurons cultures at high cell density. Cells were lysed for 20 minutes at 4 °C in RIPA-buffer (25 mM Tris-HCl pH 7.6, 150 mM NaCl, 1% NP-40, 1% sodium deoxycholate, 0.1% SDS) supplemented with protease inhibitors (Sigma). Protein extracts were centrifuged for 15 minutes at 13,000 rpm at 4 °C. Supernatants were considered as total extracts. Protein concentration was quantified by the BCA method (Bio-Rad).

### Cross-linking and immunoprecipitation

For cross-linking assays, PC12 cell extracts were incubated with the reversible homobifunctional cross-linker dithiobis(succinimidylpropionate) (DSP; Pierce), following the manufacturer’s instructions. Reaction was quenched with Tris buffer (pH 7.5), and cells were scraped in cold lysis buffer (PBS, 5 mM EDTA, 0.5% Triton X-100). Protein extracts were centrifuged for 15 minutes at 13,000 rpm at 4 °C. Supernatants were incubated with anti-Gal-4 or anti-contactin 1 antibodies at 4 °C for 18 hours, and then complexes were bound to protein-A- or protein-G-Sepharose. The immunoprecipitated complexes were separated by SDS-PAGE and subjected to detection by WB for contactin 1 or Gal-4.

### Gal-4-HA expression

HA-tagged Gal-4 expression vector was prepared by insertion of the human Gal-4 cDNA into the pcDNA3.1(+)_HA vector (Invitrogen). Astrocytes or neurons were transfected in suspension using the Nucleofector system (Amaxa) according to the manufacturer’s instructions. 48/72 hours post-nucleofection, cells were harvested and processed for biochemical analysis, or fixed for ICC and morphological evaluation.

### Myelin extracts

Myelin extracts were prepared from optic and sciatic nerves from adult rats^48^. Nerves were freed from blood vessels and connective tissue in PBS, and lysed by a glass-teflon homogeinizer in homogenization buffer (20mM Hepes-NaOH, pH 7.5) supplemented with protease inhibitors. Tissue extract was centrifuged for 3 minutes at 3,000 rpm at 4 °C. Supernatant were considered as total extracts, and was centrifuged again on a Beckmann ultracentrifuge for 20 minutes at 20,000 rpm at 4 °C. Membrane pellets were then extracted in 300 μl of buffer and, this solution then loaded on a sucrose-step gradient (300 μl of 0.8 M; 300 μl of 1 M; 100 μl of 1.2 M sucrose in buffer) and centrifuged for 1 hour at 42,000 rpm. Myelin was removed from the band above the 0.8 M sucrose, re-suspended in buffer with protease inhibitors, and centrifuged on a desktop centrifuge to remove the remaining sucrose. Pellet was re-suspended in buffer without protease inhibitors for further use and analysed by WB.

### Quantification of G4Ds

Neuron cultures or neuron/OLG co-cultures were fixed at different div and immunostained for Gal-4 in non-permeabilizing conditions. After permeabilization, axons were immunostained for Tuj1, and OLGs/myelin were labelled for MBP. Images of twenty fields per condition were obtained under identical microscope settings, and Gal-4 fluorescence intensity in neuron cultures was measured using ImageJ software. Gal-4-positive segments were manually traced, and their lengths were measured using the NeuronJ plugin for ImageJ software.

To quantify the extent of coincidence between myelinated and Gal-4-containing axon segments, Gal-4 and MBP fluorescence intensity were measured along single axons in microscope images of neuron/OLG co-cultures. To standardize measurements, the maximum intensity in each channel was considered as 100 %, and the rest of intensities were expressed as a percentage of the maximum.

### Immunohistochemistry and co-localization analyses

Postnatal 3 and 30 days old Wistar rats (P3 and P30) were deeply anesthetized and transcardially perfused with saline followed by fixative solution of 4% paraformaldehyde (PFA). Brains were dissected out, post-fixed during 2 hours at RT, washed with phosphate buffer (0.2M PB, pH 7.4), and 50 μm thick coronal tissue sections were obtained with a vibrating-blade microtome (Leica VT1000S).

For IHC analysis, free-floating sections were blocked with PBS containing 2% BSA, 8% donkey serum and 0.2% Triton X-100 (blocking solution) for 1 hour at RT, and incubated with primary antibodies o/n at 4 °C. After extensive washing, sections were incubated with secondary antibodies for 90 min, thoroughly washed, and mounted with Mowiol-DABCO. Equivalent sections of both P3 and P30 brains were selected, and mosaic images spanning the entire somatosensory cortex layers up to the striatum were acquired on a confocal microscope (Leica SP5).

Average fluorescence intensities of Gal-4 and MBP channels were measured along the entire columns from cortical layer VI to brain surface, using ImageJ software. Measurements were then expressed as percentage of the maximum fluorescence intensity in each channel, and plotted versus the distance from the bottom as expression profiles.

For co-localization analyses, Gal-4 and MBP channels from maximum projections or single z-plane confocal images were converted to gray scale images (8 bit, 72 pixels per inch), and thresholded using ImageJ software. A third particle map was obtained by applying the colocalization plugin (ImageJ) to Gal-4 and MBP images simultaneously. Number and size of Gal-4, MBP and co-localized particles were obtained using the particle analysis plugin (ImageJ). Only particles over 7 pixels were considered as positive and included in subsequent calculations.

All procedures involving animals complied with international guidelines on the ethical use of animals in the European Communities Council Directive dated 24 November 1986 (86/6091EEC) and with the guidelines of the Institutional Ethics Committee of the Hospital Nacional de Parapléjicos de Toledo (SESCAM).

## Acknowledgements

We thank Dr. Jose A. Rodriguez-Alfaro and Dr. Javier Mazario (Microscopy Facility of the Hospital Nacional de Paraplejicos –SESCAM-, Toledo, Spain) for their valuable technical support. This work was funded by i) EC Marie Curie RTN (contract no. 2005-019561), ii) EU 7^th^ Framework Program (contract no. 317297), iii) Verein zur Förderung des biologisch-technologischen Förtschritts e.V. (Heidelberg, Germany), and iii) Carlos III Health Institute and Castilla-La-Mancha Health Service (SESCAM) EMER program (EMER07/026).

## Author Contributions Statement

NDR and AMH contributed equally in designing and performing the experiments they analysed data, prepared the figures, and wrote the Methods section. They also discussed and revised the manuscript. SV designed, performed and analysed the data from oligodendrocyte and co-culture experiments. MPI participated in cell culture preparation and maintenance, and performed the histochemical experiments. HJG produced and provided recombinant galectins, antibodies and galectin-related genetic material. He actively participated in the analysis and discussion of results and revised the manuscript. JAR conceived the study and designed investigations, participated in some experiments and data analysis, and wrote the manuscript. All authors read and approved the final manuscript.

## Additional Information

Competing Financial Interests: The authors declare that they have no competing interests.

